# An Oxidative Stress-Associated Seven-Gene Prognostic Signature in Lung Adenocarcinoma: Integrative Transcriptomic Analysis Across Public Cohorts

**DOI:** 10.64898/2026.09.04.749312

**Authors:** Xingchen Zhou, Zhen Le, Pengxia Song, Qianru Xu, Ming Chen, Xiaobo Liu, Sijie Zhan, Yubin Liu, Lin Zhang

## Abstract

Lung adenocarcinoma is molecularly heterogeneous, and oxidative-stress programs can support either tumor restraint or tumor adaptation depending on cellular context. This study integrated public lung adenocarcinoma transcriptomic cohorts to identify oxidative-stress-associated expression features and evaluate their prognostic relevance. Expression profiles from The Cancer Genome Atlas, Genotype-Tissue Expression project, and GEO series GSE31210, GSE40791, and GSE30219 were analyzed. Differential expression, weighted gene co-expression network analysis, functional enrichment, univariable Cox regression, and least absolute shrinkage and selection operator Cox modeling were combined to derive a risk signature. Immune-cell enrichment, gene set enrichment analysis, gene set variation analysis, and pan-cancer analyses were used for biological characterization. A total of 1,305 genes differed between tumor and control samples, including 498 upregulated and 807 downregulated genes. Intersection of differentially expressed genes, the oxidative-stress-associated co-expression module, and the oxidative-stress gene set yielded 44 genes enriched in responses to reactive oxygen species and hydrogen peroxide, antioxidant and peroxidase activities, focal adhesion, Rap1 signaling, and PI3K-Akt signaling. A seven-gene signature comprising *FBLN5, HBB, FYN, HGF, TFAP2A, PLIN5*, and *F2RL1* stratified the 523-sample training cohort and the 207-sample internal validation cohort into groups with different overall survival. Time-dependent areas under the receiver operating characteristic curve at 1, 3, and 5 years were 0.677, 0.622, and 0.649 in training and 0.613, 0.691, and 0.706 in internal validation. In the 85-case GSE30219 external cohort, corresponding values were 0.588, 0.661, and 0.631; survival separation followed the expected direction but did not reach statistical significance (log-rank P = 0.100). Seventeen immune-cell signatures differed between risk groups, while high-risk tumors were enriched for cell-cycle, DNA-replication, mismatch-repair, glycolytic, E2F, G2M-checkpoint, MYC-target, and mTORC1-related programs. The signature therefore captures reproducible oxidative-stress-associated transcriptional variation with moderate prognostic discrimination. Its clinical utility requires prospective evaluation, complete clinical adjustment, and experimental validation.

## 1. Introduction

Lung cancer remains a leading cause of cancer mortality worldwide, and adenocarcinoma is its most common histological subtype [1]. Large-scale genomic studies have established that lung adenocarcinoma comprises biologically distinct tumors with diverse driver alterations, transcriptional states, and clinical trajectories [2]. This heterogeneity limits the prognostic resolution of conventional clinicopathological variables and motivates the development of molecular models that describe disease-relevant processes rather than isolated markers. Transcriptome-based models can quantify such processes across large retrospective cohorts, provided that model development is coupled to independent validation and that predictive performance is interpreted within the limitations of the available data.

Reactive oxygen species are integral components of cellular signaling and homeostasis, but excessive or inadequately buffered oxidant production damages nucleic acids, proteins, and lipids [3]. Cancer cells are exposed to oxidative pressure generated by oncogenic signaling, altered metabolism, mitochondrial dysfunction, hypoxia, inflammation, and treatment. At the same time, tumors can increase antioxidant capacity and use redox signaling to support proliferation, survival, invasion, and therapy resistance [4]. Oxidative stress is therefore not a unidirectional tumor-promoting or tumor-suppressive process. Its biological effects depend on ROS abundance, subcellular source, antioxidant buffering, tissue compartment, and disease stage, consistent with the broader concept that cancer phenotypes arise from interacting cellular and microenvironmental programs [5].

Public transcriptomic resources enable the study of these programs at cohort scale. The Cancer Genome Atlas (TCGA), the Genotype-Tissue Expression (GTEx) project, and the Gene Expression Omnibus (GEO) provide complementary tumor, adjacent-tissue, normal-tissue, and outcome data [2,6,7]. Network approaches such as weighted gene co-expression network analysis can identify groups of coordinately expressed genes linked to a phenotype, while penalized survival regression can reduce correlated candidates to a parsimonious model [18,20]. However, risk signatures derived from retrospective public data frequently show attenuated performance during external testing, and their biological interpretation must distinguish expression associations from experimentally demonstrated mechanisms.

The present study integrated TCGA, GTEx, and three GEO lung adenocarcinoma cohorts to define oxidative-stress-associated transcriptional features. Differential expression and co-expression network analysis were combined with an oxidative-stress gene set to nominate candidate genes. A seven-gene Cox model was then evaluated in training, internal validation, and external validation cohorts. Immune-cell enrichment, pathway-level analyses, and pan-cancer comparisons were used to characterize the molecular context captured by the risk score. The analysis was designed to identify a reproducible prognostic signal while maintaining a conservative boundary between computational association and biological causation.

## 2. Materials and Methods

### 2.1 Study design and public transcriptomic cohorts

This retrospective computational study used publicly available expression and clinical data from TCGA, GTEx, and GEO. TCGA-LUAD and GTEx lung data supplied tumor and normal-tissue expression profiles, and GEO series GSE31210, GSE40791, and GSE30219 supplied independent microarray cohorts generated on the Affymetrix Human Genome U133 Plus 2.0 Array platform (GPL570). The processed expression matrices and accompanying phenotype data were downloaded from their source repositories and underwent quality control and normalization before analysis. The analysis record designated TCGA-LUAD and GSE31210 as derivation data, GSE40791 as an external tumor-versus-normal expression cohort, and the 85-case lung adenocarcinoma survival subset of GSE30219 as an external prognostic-validation cohort. The original studies describing GSE31210, GSE30219, and GSE40791 are cited in references [9–11].

### 2.2 Oxidative-stress gene collection

Oxidative-stress-associated genes were obtained from the Molecular Signatures Database (MSigDB) [12]. The initial collection contained 467 genes. Differential-expression filtering retained 61 oxidative-stress-associated genes for network intersection and downstream analyses.

### 2.3 Differential-expression analysis

Differential expression between lung adenocarcinoma and control samples was assessed with the limma package (version 3.50.0) [13]. Genes with an absolute log2 fold change greater than 1 and a Benjamini-Hochberg-adjusted P value below 0.05 were considered differentially expressed [14]. The analysis summarized the numbers of upregulated and downregulated genes and visualized the results by volcano plot, expression heatmap, and gene-level boxplots.

### 2.4 Functional enrichment analysis

Gene Ontology (GO) biological process, cellular component, and molecular function terms and Kyoto Encyclopedia of Genes and Genomes (KEGG) pathways were evaluated for the full differential-expression set and for the oxidative-stress-associated intersection [15,16]. Enrichment analysis was performed with clusterProfiler (version 4.2.2) [17], using P < 0.05 as the reported significance threshold.

### 2.5 Weighted gene co-expression network analysis

WGCNA (version 1.70-3) was applied to a standardized expression matrix containing 739 lung adenocarcinoma tumors [18]. Pearson correlations were transformed into a weighted adjacency matrix, and a soft-thresholding power of beta = 7 was selected to approximate scale-free topology while preserving network connectivity. The adjacency matrix was converted to a topological overlap matrix, hierarchical clustering and dynamic tree cutting were used to define modules, and unassigned genes were placed in the grey module. Module eigengenes were correlated with the oxidative-stress phenotype. The module with the strongest association was selected, and the relationship between module membership and gene significance was evaluated by Pearson correlation.

### 2.6 Identification of oxidative-stress-associated lung adenocarcinoma genes

Candidate genes were defined by intersecting three sets: genes differentially expressed between lung adenocarcinoma and control samples, genes assigned to the oxidative-stress-associated WGCNA module, and the filtered oxidative-stress gene collection. The resulting intersection was analyzed by GO and KEGG enrichment and was carried forward to survival screening.

### 2.7 Construction and validation of the prognostic model

Associations between candidate-gene expression and overall survival were first screened by univariable Cox proportional-hazards regression [19]. Genes with P <= 0.05 were entered into least absolute shrinkage and selection operator Cox regression implemented with glmnet [20]. The reported complete-case prognostic dataset contained 730 tumors and was randomly divided in a 7:3 ratio into a training cohort of 523 cases and an internal validation cohort of 207 cases, using random seed 8655. The penalty parameter was selected at the minimum cross-validated partial-likelihood deviance. For each patient, the risk score was calculated as the sum of the standardized expression value of each selected gene multiplied by its fitted LASSO-Cox coefficient. Patients were separated into high-risk and low-risk groups using the cohort-specific median risk score.

Overall-survival distributions were estimated with the Kaplan-Meier method and compared by the log-rank test [21]. Time-dependent receiver operating characteristic analysis quantified discrimination at 1, 3, and 5 years [22]. The fitted seven-gene model was evaluated in the internal validation cohort and in the independent 85-case GSE30219 lung adenocarcinoma survival cohort. Risk-score distributions, survival status, model-gene expression, Kaplan-Meier curves, and time-dependent receiver operating characteristic curves were visualized for each cohort.

### 2.8 Immune-cell enrichment analysis

Gene signatures representing 28 immune-cell populations were obtained from TISIDB [23]. Single-sample gene-set enrichment scores were calculated from tumor expression profiles to estimate the relative enrichment of each immune-cell signature. Scores were compared between high-risk and low-risk groups, and pairwise correlations among immune-cell signatures were calculated. Plots were generated with ggplot2 (version 3.3.6) [26].

### 2.9 GSEA and GSVA

For gene set enrichment analysis (GSEA), all genes were ranked by log2 fold change between high-risk and low-risk tumors [24]. clusterProfiler (version 4.2.2) was used with 1,000 gene-set permutations and the c2.cp.kegg.v7.5.1.symbols collection from MSigDB. Gene sets with P < 0.05 were reported as significantly enriched. Gene set variation analysis (GSVA; version 1.42.0) was performed with MSigDB hallmark gene sets to compare pathway-level activity across individual tumors [25]. The resulting matrix was visualized with pheatmap (version 1.0.12).

### 2.10 Pan-cancer expression, survival, and immune-correlation analysis

Expression and clinical data for 33 TCGA cancer types were retrieved with TCGAbiolinks (version 2.25.0) [8]. For model genes represented in the pan-cancer output, tumor-normal expression differences were tested by the Wilcoxon rank-sum test. Univariable Cox regression estimated associations with overall survival, and single-sample gene-set enrichment scores were used to examine correlations between gene expression and immune-cell signatures across cancer types.

### 2.11 Statistical analysis

Continuous variables were compared between two groups with the nonparametric Wilcoxon rank-sum test. Categorical proportions were evaluated with the chi-square test or Fisher exact test, as appropriate. Kaplan-Meier curves were fitted with survminer, and survival differences were assessed by two-sided log-rank tests. Two-sided P < 0.05 was considered statistically significant unless an adjusted threshold was specified. All analyses were performed in R version 4.1.2 [27].

## 3. Results

### 3.1 Lung adenocarcinoma shows extensive transcriptomic differences from control lung tissue

Differential-expression analysis identified 1,305 genes that met the prespecified fold-change and significance criteria. Among these genes, 498 were upregulated and 807 were downregulated in lung adenocarcinoma relative to control tissue (Figure 1A). The most strongly upregulated genes displayed in the heatmap were *OCIAD2, GOLM1, PYCR1, ALDH18A1*, and *B3GNT3*, whereas *FENDRR, STARD8, LIMS2, TAL1*, and *ACVRL1* were among the most strongly downregulated genes (Figure 1B). Gene-level boxplots confirmed pronounced tumor-control separation for the 20 displayed genes (Figure 1C).

**Figure 1.**
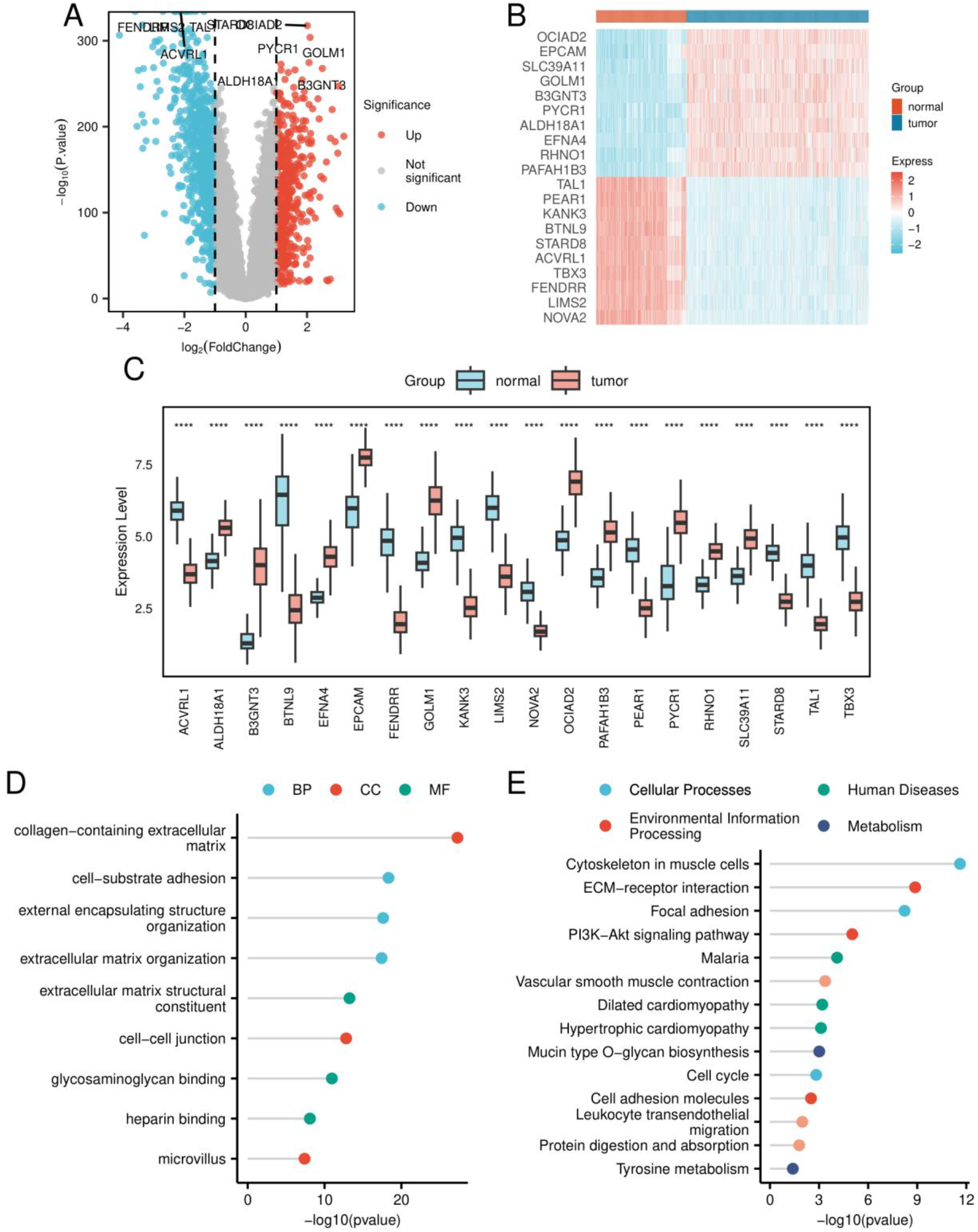
Differentially expressed genes between lung adenocarcinoma and control lung tissue. (A) Volcano plot of the tumor-control comparison. Red points denote upregulated genes, blue points denote downregulated genes, and grey points denote genes outside the differential-expression criteria. (B) Heatmap of the most strongly differentially expressed genes across tumor and control samples. (C) Boxplots of 20 representative differentially expressed genes. (D) Representative Gene Ontology enrichment terms, grouped as biological process, cellular component, and molecular function. (E) Representative KEGG pathways, grouped by KEGG functional category. Asterisks in panel C indicate significance: ****P < 0.0001, ***P < 0.001, **P < 0.01, and *P < 0.05.

The differential-expression set was enriched for extracellular-matrix organization, cell-substrate adhesion, external encapsulating-structure organization, collagen-containing extracellular matrix, cell-cell junctions, extracellular-matrix structural constituents, glycosaminoglycan binding, and heparin binding (Figure 1D). KEGG analysis further highlighted cytoskeletal organization, focal adhesion, ECM-receptor interaction, PI3K-Akt signaling, cell-cycle regulation, cell-adhesion molecules, leukocyte transendothelial migration, mucin-type O-glycan biosynthesis, protein digestion and absorption, and amino-acid metabolic pathways (Figure 1E). Together, these results indicated that the tumor-control contrast involved coordinated changes in extracellular architecture, adhesion, proliferation, and metabolism rather than a single isolated pathway.

### 3.2 Co-expression network analysis identifies an oxidative-stress-associated module

The soft-thresholding analysis supported beta = 7, at which the scale-free topology fit exceeded 0.85 and mean connectivity approached zero (Figure 2A). Hierarchical clustering and dynamic tree cutting identified nine non-grey co-expression modules (Figure 2B). The eigengene-adjacency heatmap showed distinct but partially related module-level expression patterns (Figure 2C). Among the modules, the turquoise module had the strongest positive association with the oxidative-stress phenotype (r = 0.8709; Figure 2D). Within the turquoise module, module membership was strongly correlated with oxidative-stress gene significance (r = 0.93, P < 1 x 10⋀-200; Figure 2E), indicating that genes most central to this module also carried the strongest oxidative-stress association.

**Figure 2.**
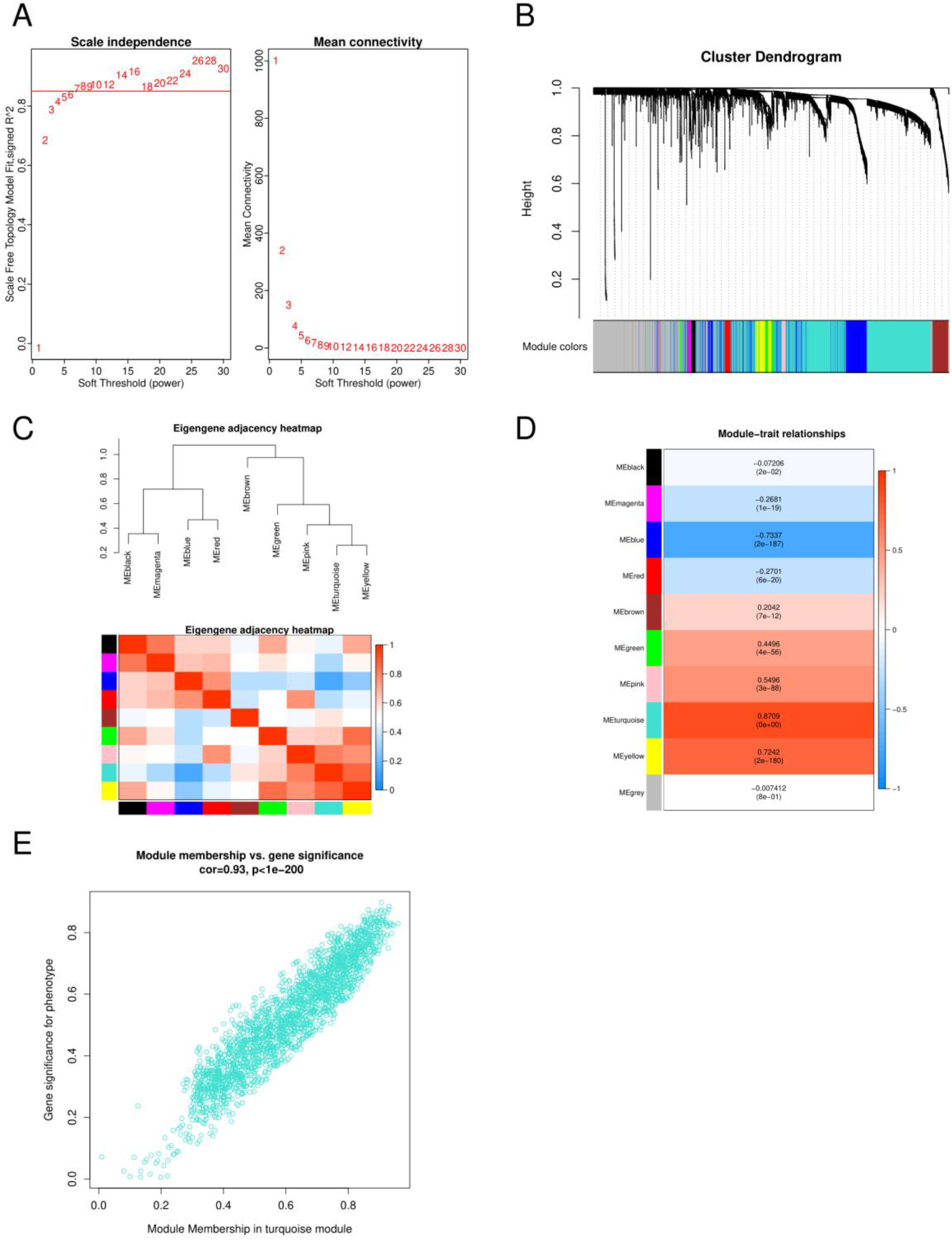
Weighted gene co-expression network analysis of oxidative-stress-associated expression patterns. (A) Scale-independence and mean-connectivity plots used to select a soft-thresholding power of beta = 7. (B) Gene-clustering dendrogram with dynamic tree-cut module assignments. (C) Eigengene-adjacency heatmap summarizing relationships among modules; red indicates higher adjacency and blue indicates lower adjacency. (D) Correlations between module eigengenes and the oxidative-stress phenotype, with correlation coefficients and corresponding P values shown in each cell. (E) Correlation between module membership and oxidative-stress gene significance for genes in the turquoise module.

### 3.3 Forty-four genes connect differential expression, co-expression, and oxidative-stress annotation

Intersection of the lung adenocarcinoma differential-expression set, the turquoise-module gene set, and the filtered oxidative-stress gene collection yielded 44 genes (Figure 3A). These genes were enriched for responses to reactive oxygen species, oxidative stress, and hydrogen peroxide; antioxidant activity; oxidoreductase activity acting on peroxide; and peroxidase activity (Figure 3C). Cellular-component terms included hemoglobin complex, haptoglobin-hemoglobin complex, and endocytic-vesicle lumen. KEGG enrichment included focal adhesion, Rap1 signaling, PI3K-Akt signaling, phospholipase D signaling, *EGFR* tyrosine-kinase-inhibitor resistance, glycosaminoglycan degradation, and infection-related pathways that share host-response genes (Figure 3D).

**Figure 3.**
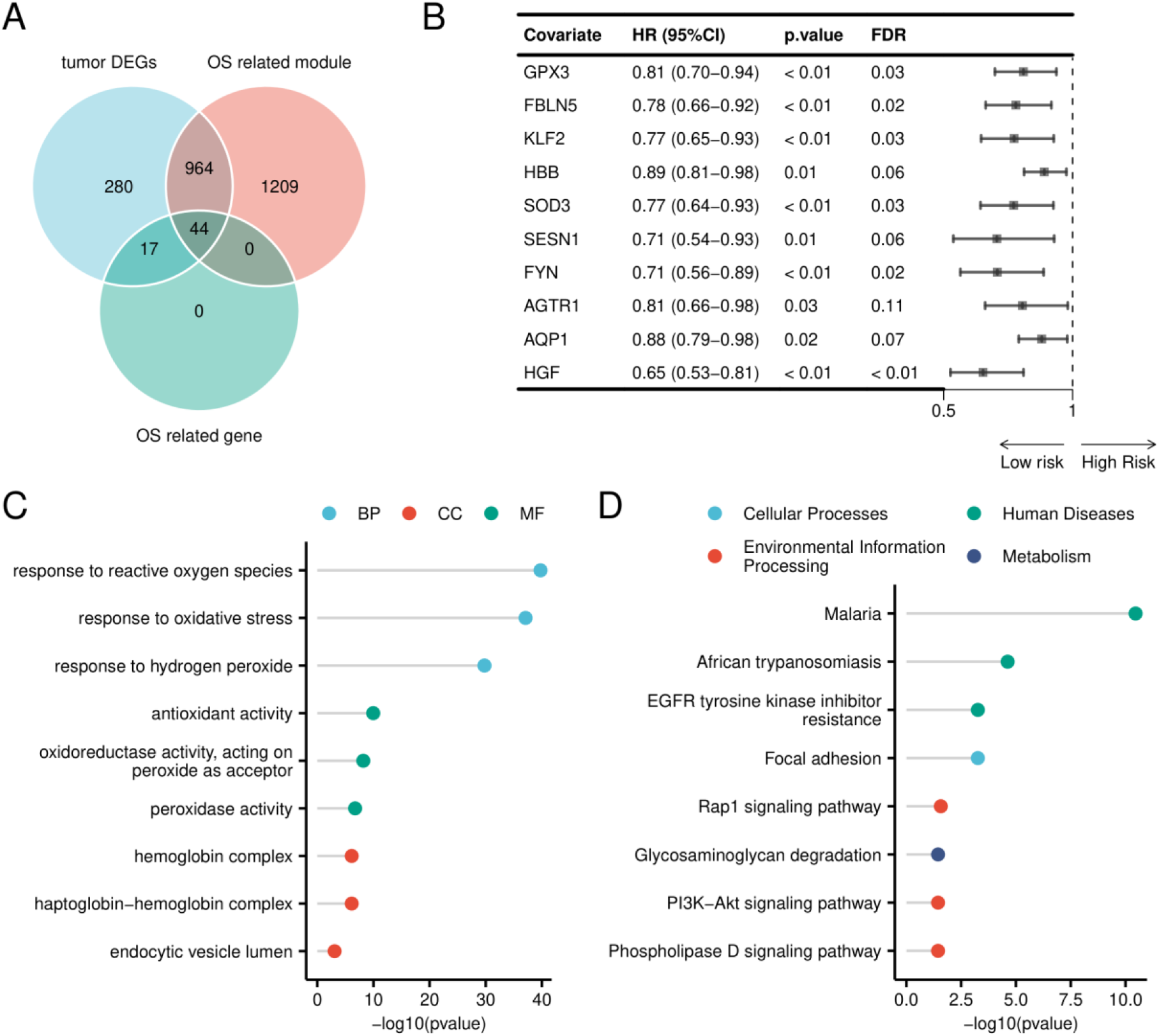
Identification and functional characterization of oxidative-stress-associated lung adenocarcinoma genes. (A) Venn diagram showing the 44-gene intersection among tumor differentially expressed genes, genes in the oxidative-stress-associated WGCNA module, and oxidative-stress-related genes. (B) Univariable Cox forest plot for ten survival-associated genes displayed in the analysis output; hazard ratios, 95% confidence intervals, nominal P values, and false-discovery-rate values are shown. (C) Representative Gene Ontology enrichment terms for the 44 genes. (D) Representative KEGG pathways enriched among the 44 genes.

Univariable Cox analysis identified survival-associated genes within the 44-gene set (Figure 3B). The displayed forest plot included *GPX3, FBLN5, KLF2, HBB, SOD3, SESN1, FYN, AGTR1, AQP1*, and *HGF*. All displayed hazard ratios were below 1.0, although the false-discovery-rate values for *HBB, SESN1, AGTR1*, and *AQP1* exceeded 0.05. These survival-screening results supplied candidates for penalized Cox modeling rather than independent evidence of clinical utility.

### 3.4 A seven-gene score stratifies survival in the training and internal validation cohorts

LASSO-Cox regression reduced the survival-screened candidates to seven genes: *FBLN5, HBB, FYN, HGF, TFAP2A, PLIN5*, and *F2RL1* (Figure 4A-B). The 523-case training cohort was divided at the median score into 261 high-risk and 262 low-risk cases. Risk-score and survival-status plots showed a higher concentration of deaths at increasing risk scores, and the expression heatmap demonstrated coordinated variation across the seven model genes (Figure 4C). High-risk patients had shorter overall survival by log-rank testing (P < 0.05; Figure 4D). The 1-, 3-, and 5-year time-dependent AUCs were 0.677, 0.622, and 0.649, respectively (Figure 4E and Table 2).

**Table 1.** Public transcriptomic cohorts and their roles in the analysis.

| Cohort | Repository / platform | Samples reported in the analysis record | Primary role |
| --- | --- | --- | --- |
| TCGA-LUAD | TCGA; RNA-expression data | 513 tumors; 59 adjacent normal tissues | Derivation expression data, survival analysis, and pan-cancer framework |
| GTEx lung | GTEx; RNA-expression data | 288 normal lung tissues | Normal-tissue reference for tumor-control comparisons |
| GSE31210 | GEO; GPL570 microarray | 226 tumors; 20 normal tissues; 226 tumors with survival information | Derivation expression and survival data; component of the 739-tumor WGCNA matrix |
| GSE40791 | GEO; GPL570 microarray | 94 tumors; 100 normal tissues | External validation of tumor-versus-normal expression patterns |
| GSE30219 | GEO; GPL570 microarray | 85 lung adenocarcinoma cases used for survival validation | External prognostic validation |

Abbreviations: GEO, Gene Expression Omnibus; GTEx, Genotype-Tissue Expression; LUAD, lung adenocarcinoma; TCGA, The Cancer Genome Atlas; WGCNA, weighted gene co-expression network analysis. Sample counts reproduce the analysis record and refer to the subsets used in the reported workflow.

**Table 2.** Prognostic performance of the seven-gene risk score.

| Cohort | Sample size | Risk-group survival comparison | 1-year AUC | 3-year AUC | 5-year AUC |
| --- | --- | --- | --- | --- | --- |
| Training | 523 | Log-rank P < 0.05 | 0.677 | 0.622 | 0.649 |
| Internal validation | 207 | Log-rank P < 0.05 | 0.613 | 0.691 | 0.706 |
| GSE30219 external validation | 85 | Log-rank P = 0.100 | 0.588 | 0.661 | 0.631 |

AUC values are time-dependent areas under the receiver operating characteristic curve. Confidence intervals were not available in the analysis output.

**Figure 4.**
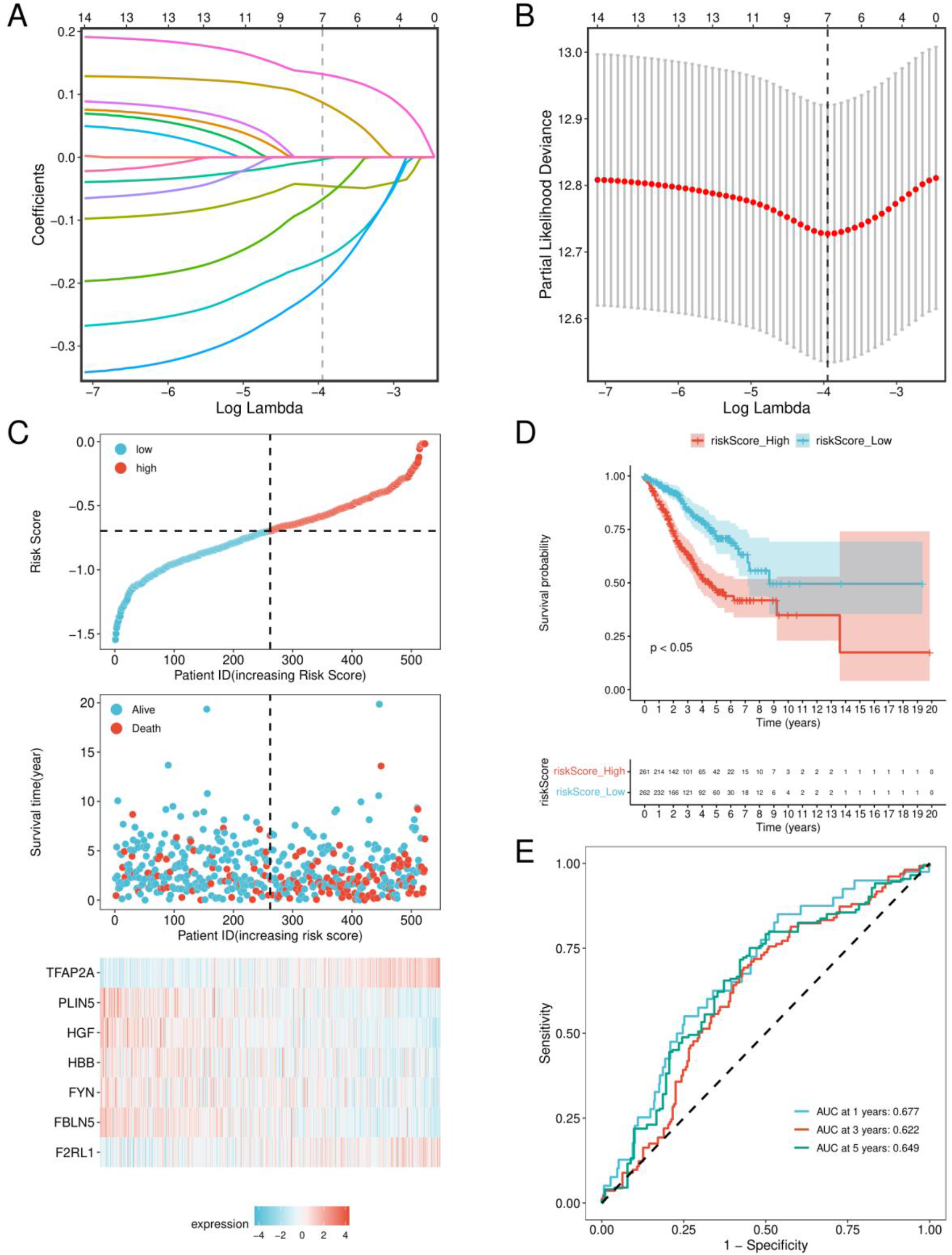
Construction of the seven-gene prognostic model in the training cohort. (A) LASSO-Cox coefficient trajectories across values of log lambda. (B) Cross-validated partial-likelihood deviance used to select the penalty parameter. (C) Risk-score distribution, survival status, and standardized expression of the seven model genes in the 523-case training cohort. (D) Kaplan-Meier overall-survival curves for the high-risk and low-risk groups. (E) Time-dependent receiver operating characteristic curves at 1, 3, and 5 years.

Application of the same seven-gene model to the 207-case internal validation cohort produced 103 high-risk and 104 low-risk cases. The risk-score distribution again aligned with a higher frequency of deaths at higher scores (Figure 5A), and the high-risk group had shorter overall survival (log-rank P < 0.05; Figure 5B). The 1-, 3-, and 5-year AUCs were 0.613, 0.691, and 0.706 (Figure 5C and Table 2). Expression comparisons showed higher *F2RL1* and *TFAP2A* in the high-risk group, whereas *FBLN5, FYN, HBB, HGF*, and *PLIN5* were lower in the high-risk group (Figure 5D).

**Figure 5.**
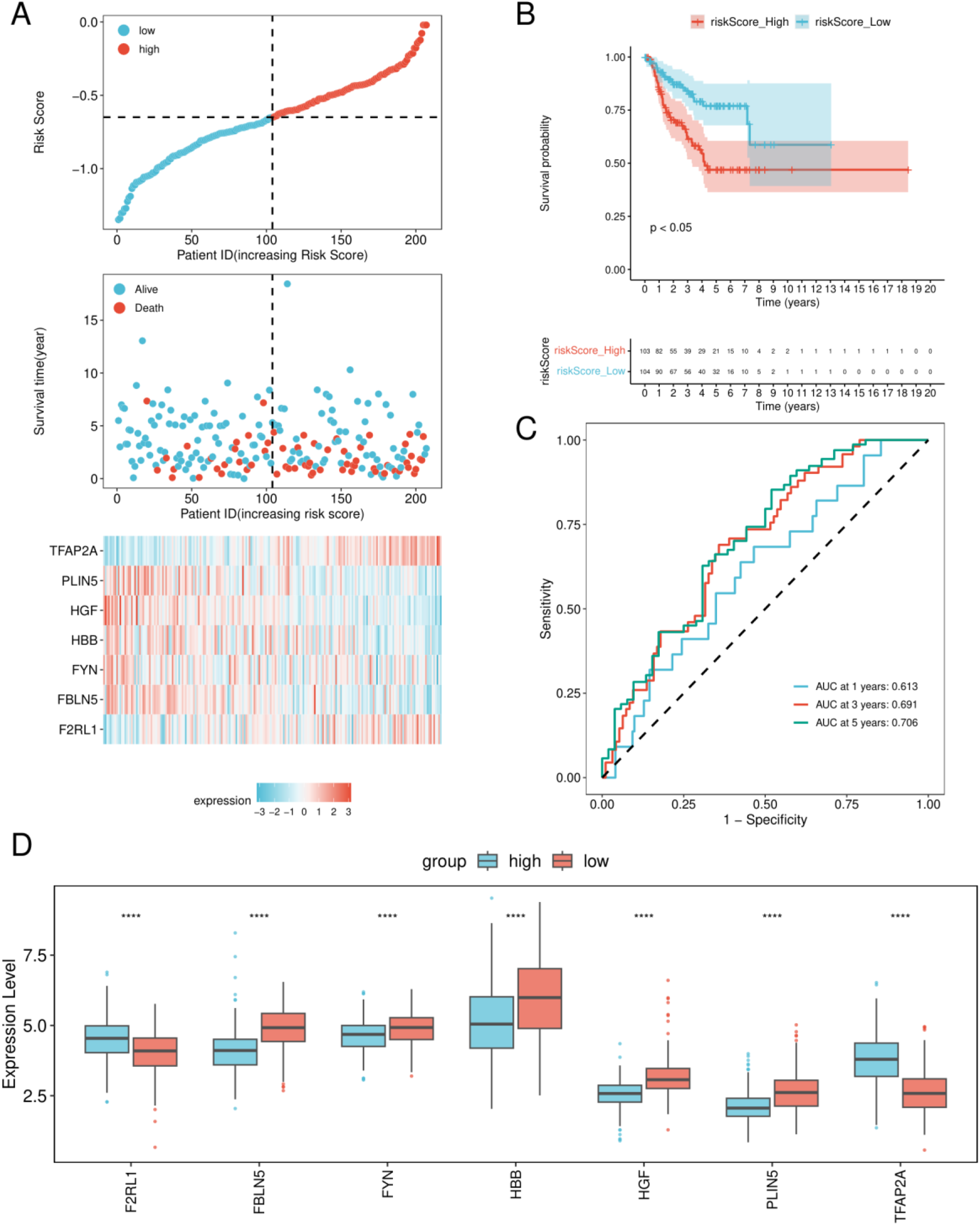
Internal validation of the seven-gene prognostic model. (A) Risk-score distribution, survival status, and standardized model-gene expression in the 207-case internal validation cohort. (B) Kaplan-Meier overall-survival curves for high-risk and low-risk patients. (C) Time-dependent receiver operating characteristic curves at 1, 3, and 5 years. (D) Expression distributions of the seven model genes by risk group. Asterisks indicate significance: ****P < 0.0001, ***P < 0.001, **P < 0.01, and *P < 0.05.

### 3.5 External validation shows moderate discrimination with nonsignificant survival separation

The independent GSE30219 lung adenocarcinoma subset contained 85 cases, divided into 42 high-risk and 43 low-risk cases. The risk distribution and survival-status plot preserved the expected increase in adverse outcomes across the score range (Figure 6A). The high-risk curve showed poorer overall survival, but the difference did not reach the prespecified significance threshold (log-rank P = 0.100; Figure 6B). External time-dependent AUCs were 0.588 at 1 year, 0.661 at 3 years, and 0.631 at 5 years (Figure 6C and Table 2), indicating moderate but attenuated discrimination relative to the internal validation cohort.

**Figure 6.**
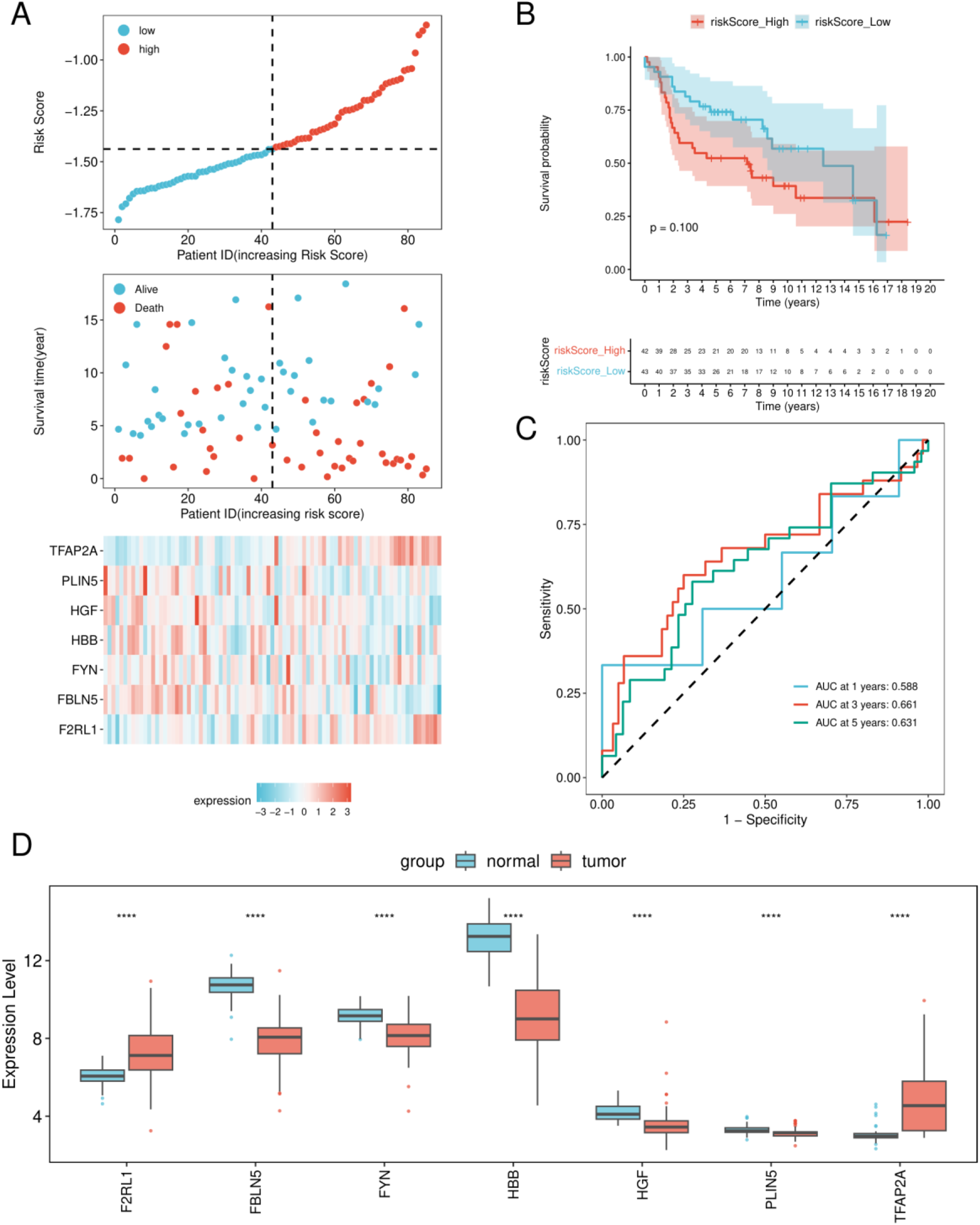
External validation of the seven-gene signature. (A) Risk-score distribution, survival status, and standardized model-gene expression in the 85-case GSE30219 lung adenocarcinoma survival cohort. (B) Kaplan-Meier overall-survival curves for high-risk and low-risk patients; log-rank P = 0.100. (C) Time-dependent receiver operating characteristic curves at 1, 3, and 5 years. (D) Independent tumor-versus-normal expression comparison for the seven model genes. In panel D, ****P < 0.0001.

An independent tumor-versus-normal expression comparison reproduced the model-gene directions observed in the derivation data: *F2RL1* and *TFAP2A* were higher in tumor tissue, whereas *FBLN5, FYN, HBB, HGF*, and *PLIN5* were lower in tumor tissue (all displayed comparisons P < 0.0001; Figure 6D). This expression-level replication supports the stability of the underlying tumor-control contrast, while the external survival result defines a more conservative estimate of prognostic generalizability.

### 3.6 Risk groups differ in immune-cell enrichment profiles

Single-sample enrichment analysis quantified 28 immune-cell signatures across the risk groups (Figure 7A). Seventeen signatures differed at P < 0.05: central memory CD8 T cells, activated CD4 T cells, T follicular helper cells, gamma-delta T cells, type 17 T helper cells, type 2 T helper cells, activated B cells, immature B cells, memory B cells, CD56bright natural killer cells, CD56dim natural killer cells, myeloid-derived suppressor cells, plasmacytoid dendritic cells, macrophages, eosinophils, mast cells, and neutrophils (Figure 7C). High-risk tumors showed higher scores for activated CD4 T cells, central memory CD8 T cells, gamma-delta T cells, T follicular helper cells, type 17 and type 2 T helper cells, memory B cells, both natural-killer-cell subsets, and plasmacytoid dendritic cells. Low-risk tumors showed higher activated and immature B-cell, macrophage, eosinophil, mast-cell, neutrophil, and myeloid-derived suppressor-cell scores.

**Figure 7.**
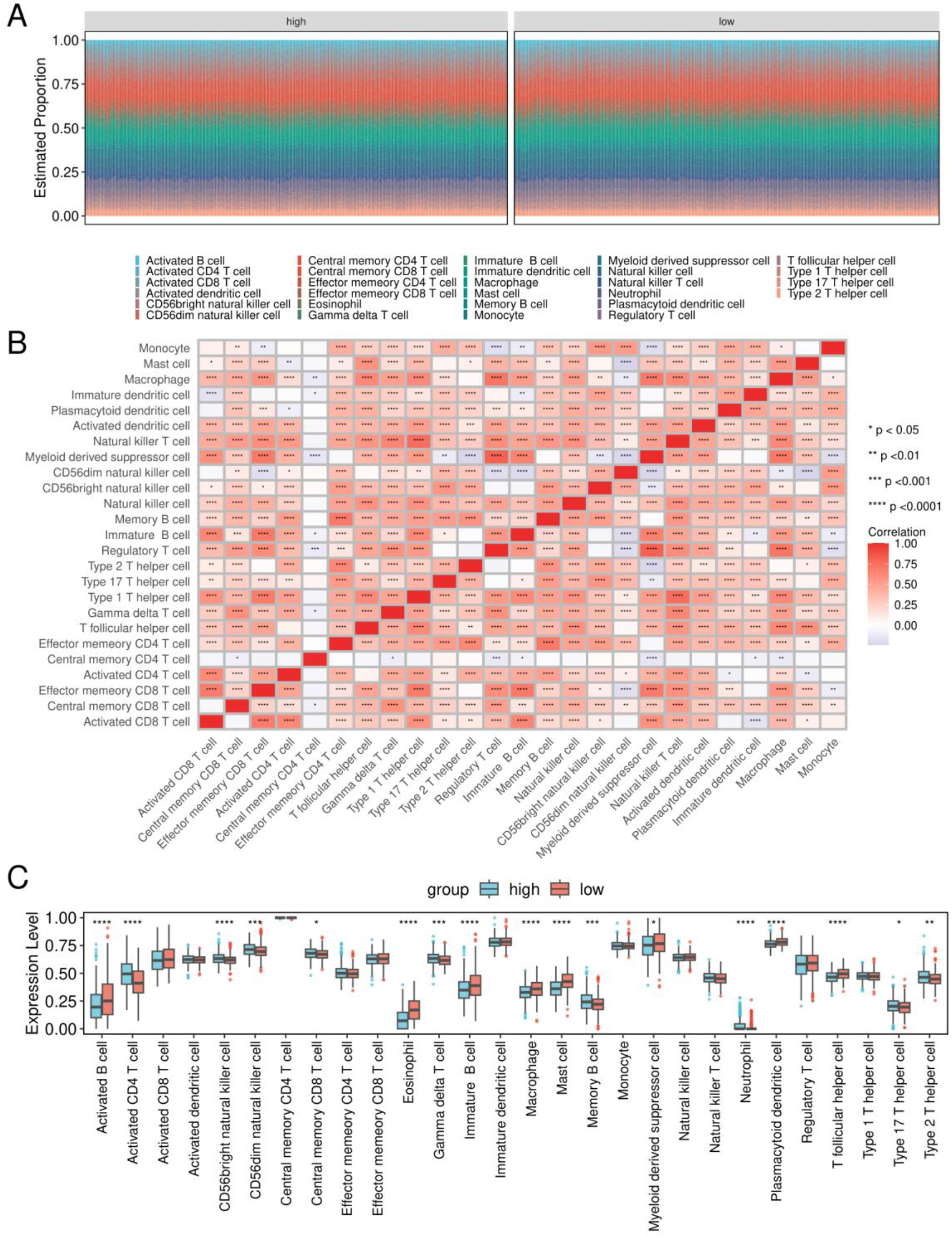
Immune-cell enrichment patterns in lung adenocarcinoma risk groups. (A) Stacked visualization of relative enrichment scores for 28 immune-cell signatures across high-risk and low-risk tumors. (B) Pairwise correlation matrix among immune-cell enrichment scores. (C) Boxplots comparing immune-cell enrichment scores between risk groups. Blue denotes high risk and red denotes low risk. Asterisks indicate significance: ****P < 0.0001, ***P < 0.001, **P < 0.01, and *P < 0.05. The plotted values are expression-derived enrichment scores and should not be interpreted as directly measured cell fractions.

The immune-cell correlation matrix contained predominantly positive relationships, with the strongest correlations concentrated among related adaptive and innate immune populations (Figure 7B). These results indicate that the risk score is associated with a coordinated shift in immune-cell enrichment rather than a uniform increase or decrease in total immune signal.

### 3.7 High-risk tumors are enriched for proliferative and biosynthetic programs

GSEA identified strong positive enrichment of cell cycle (normalized enrichment score [NES] = 2.59), DNA replication (NES = 2.51), and mismatch repair (NES = 2.18) toward the high-risk end of the ranked expression profile (Figure 8A-C and Table 3). Vascular smooth muscle contraction (NES = −2.18), systemic lupus erythematosus (NES = −2.17), and asthma (NES = −2.14) were enriched toward the low-risk end (Figure 8D-F). The negative pathways include immune, contractile, and tissue-state gene components and therefore represent coordinated expression patterns rather than diagnoses in the analyzed patients.

**Table 3.**
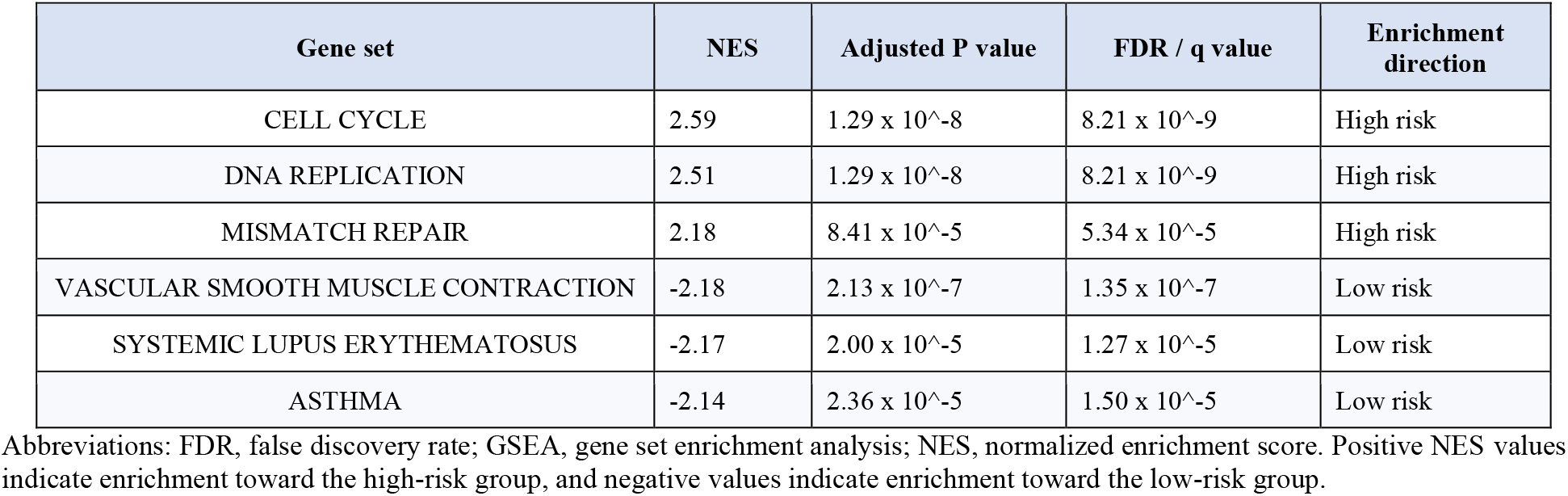
Representative GSEA results for high-risk versus low-risk lung adenocarcinoma.

**Figure 8.**
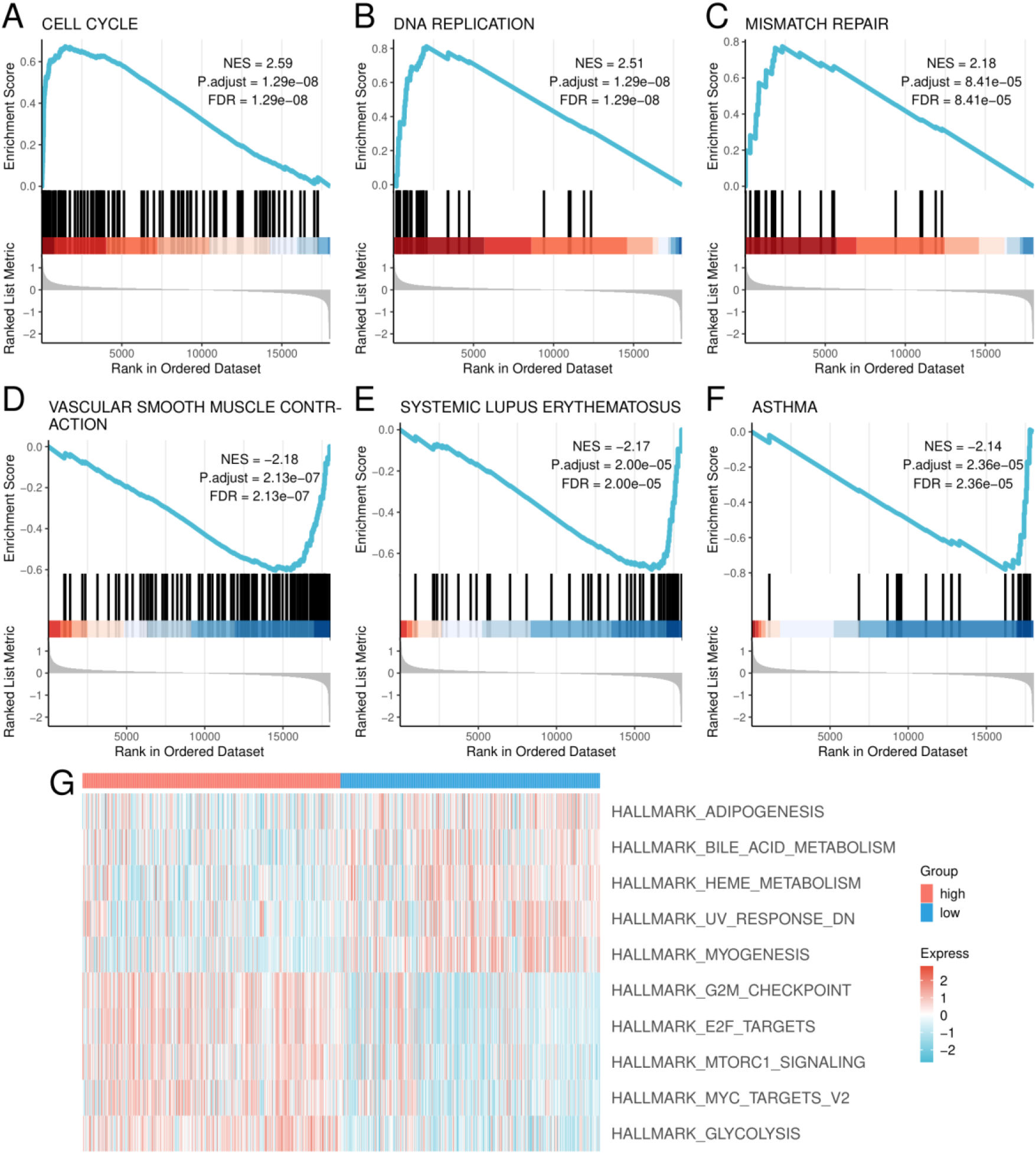
Pathway differences between high-risk and low-risk lung adenocarcinoma. (A-F) GSEA enrichment plots for cell cycle, DNA replication, mismatch repair, vascular smooth muscle contraction, systemic lupus erythematosus, and asthma. Each panel reports the normalized enrichment score, adjusted P value, and false-discovery-rate value. (G) GSVA heatmap of representative hallmark pathways differing between risk groups. Red indicates higher standardized pathway scores and blue indicates lower scores.

GSVA using hallmark gene sets identified 13 programs with higher scores in the high-risk group, including glycolysis, E2F targets, G2M checkpoint, MYC targets V2, and mTORC1 signaling. Nineteen programs had lower scores in the high-risk group, including myogenesis, heme metabolism, ultraviolet response down, bile-acid metabolism, and adipogenesis (Figure 8G). The combined results link the risk score to increased proliferation and biosynthetic activity and to reduced expression of differentiated metabolic and tissue-structural programs.

### 3.8 Model genes show context-dependent pan-cancer associations

Pan-cancer analyses were available for *F2RL1, FBLN5, FYN, HGF, PLIN5*, and *TFAP2A* and are presented in Supplementary Figures S1-S6. Each output combined univariable Cox regression across TCGA cancers, tumor-normal expression comparisons, and correlations with 28 immune-cell signatures. The direction and magnitude of associations varied across cancer types, demonstrating that the model genes do not behave as universal risk or protective factors.

Within TCGA-LUAD, higher *FBLN5, FYN, HGF*, and *PLIN5* expression was associated with lower hazard, whereas higher *TFAP2A* and *F2RL1* expression was associated with higher hazard (Table 4). These transcript-level associations were concordant with the directions of the seven-gene risk model. Immune correlations also differed across tumor types, supporting a context-dependent relationship between model-gene expression and the tumor microenvironment.

**Table 4.** Univariable overall-survival associations for six model genes in TCGA-LUAD pan-cancer output.

| Gene | Hazard ratio | 95% confidence interval | P value | Association direction |
| --- | --- | --- | --- | --- |
| FBLN5 | 0.64 | 0.46-0.89 | 0.008 | Lower hazard with higher expression |
| FYN | 0.64 | 0.47-0.87 | 0.005 | Lower hazard with higher expression |
| HGF | 0.67 | 0.48-0.93 | 0.016 | Lower hazard with higher expression |
| PLIN5 | 0.55 | 0.41-0.75 | <0.001 | Lower hazard with higher expression |
| TFAP2A | 2.04 | 1.51-2.75 | <0.001 | Higher hazard with higher expression |
| F2RL1 | 1.80 | 1.33-2.44 | <0.001 | Higher hazard with higher expression |

The values were read from the TCGA-LUAD rows of the corresponding pan-cancer forest plots. A pan-cancer output for *HBB* was not present in the source analysis.

## 4. Discussion

This study integrated differential expression, co-expression network structure, survival modeling, immune-cell enrichment, and pathway analysis to examine oxidative-stress-associated transcriptional variation in lung adenocarcinoma. The analysis identified 1,305 tumor-control differentially expressed genes, a strongly oxidative-stress-associated turquoise co-expression module, and a 44-gene intersection enriched for canonical redox-response functions. Penalized Cox regression yielded a seven-gene risk score that separated survival in the training and internal validation cohorts. External testing preserved moderate 3- and 5-year discrimination, but the survival comparison did not reach statistical significance. This pattern supports the existence of a reproducible expression signal while defining a realistic boundary for its present prognostic strength.

The initial tumor-control analysis was dominated by extracellular-matrix, adhesion, cytoskeletal, and PI3K-Akt-related terms. These findings are consistent with the central role of tissue architecture and cell-matrix signaling in invasion and progression [5]. The oxidative-stress intersection sharpened this broad tumor signature toward responses to reactive oxygen species and hydrogen peroxide, antioxidant and peroxidase activities, and hemoglobin-associated cellular components. The enrichment pattern indicates that redox biology in bulk lung adenocarcinoma transcriptomes includes both tumor-cell programs and contributions from stromal, vascular, erythroid, and immune compartments. Accordingly, the 44-gene set should be interpreted as an oxidative-stress-associated tissue signature rather than a tumor-cell-autonomous pathway.

Several model genes have prior biological links that are directionally consistent with the computational results. *FBLN5* is an extracellular-matrix protein that suppressed lung-cancer invasion and metastasis in experimental models by limiting *MMP7* expression [28]. Its lower expression in tumors and its hazard ratio below 1.0 in TCGA-LUAD are compatible with a protective tissue-structural role. *F2RL1* encodes protease-activated receptor 2. Experimental lung adenocarcinoma studies have linked increased *F2RL1* expression to proliferation, invasion, angiogenesis, and *EGFR*-related signaling [32,33], consistent with its higher expression in tumors, enrichment in the high-risk group, and adverse LUAD hazard ratio. *TFAP2A* has also been implicated in lung adenocarcinoma metastatic behavior [29], supporting its adverse direction in the model.

*PLIN5* links lipid-droplet biology to mitochondrial metabolism and can protect cells from oxidative injury by controlling fatty-acid handling [30]. Its lower tumor expression and protective survival association suggest that loss of differentiated lipid-storage and redox-buffering programs may accompany aggressive disease. The transcript-level *HGF* association requires more nuanced interpretation. *HGF*-*MET* signaling is a well-established mediator of lung-cancer growth and therapeutic resistance, yet *HGF* is often produced by stromal cells rather than malignant epithelial cells [31]. A protective association in bulk RNA data can therefore reflect tissue composition, tumor purity, or compartment-specific expression and does not negate the pro-tumor activity of paracrine *HGF*-*MET* signaling. A similar distinction applies to *FYN*: the current transcript-level association was protective, whereas activated phosphorylated Fyn has been associated with poorer outcome in resected lung adenocarcinoma [34]. Expression abundance and kinase activation state are not interchangeable measurements.

The pathway analyses provide a coherent systems-level interpretation of the risk score. High-risk tumors were enriched for cell cycle, DNA replication, mismatch repair, glycolysis, E2F targets, G2M checkpoint, MYC targets, and mTORC1 signaling. These programs describe a proliferative and biosynthetically active state that can increase both ROS production and dependence on antioxidant defenses [3,4]. Conversely, low-risk tumors showed greater representation of heme metabolism, myogenesis, adipogenesis, bile-acid metabolism, and vascular smooth-muscle-contraction-related genes, consistent with retention of differentiated tissue and stromal programs. The observed enrichment should not be interpreted as direct activation or inhibition of a pathway; it reflects coordinated differences in expression ranks or pathway-level scores.

Immune-cell enrichment also differed between risk groups, but the pattern was not a simple immune-hot versus immune-cold contrast. High-risk tumors had higher enrichment scores for several activated T-cell, natural-killer-cell, and plasmacytoid-dendritic-cell signatures, whereas low-risk tumors had higher scores for activated and immature B cells, macrophages, granulocytes, mast cells, and myeloid-derived suppressor cells. These shifts may reflect differences in tumor purity, inflammatory state, lymphoid organization, or compensatory immune recruitment. Because the values were inferred from bulk expression, they quantify relative signature enrichment rather than directly measured cell proportions. Single-cell or spatial profiling would be required to determine the cellular source of the redox-associated genes and the organization of the immune microenvironment.

The external validation result is central to interpretation of the model. The 3-year and 5-year AUCs of 0.661 and 0.631 indicate that the score retained some discriminatory information across datasets, but the 1-year AUC was 0.588 and the external log-rank P value was 0.100. The signature should therefore be regarded as a research model with moderate prognostic discrimination rather than a clinically established biomarker. The attenuation is compatible with cohort differences in stage composition, follow-up, platform, preprocessing, and biological heterogeneity. It also illustrates why internal validation alone can overstate model transportability.

Several limitations constrain the present analysis. First, the study was retrospective and based entirely on public bulk-transcriptomic datasets. Second, the derivation cohorts used different expression platforms, and the available analysis record does not provide enough preprocessing detail to evaluate all cross-platform effects. Third, the model was evaluated primarily by risk-group Kaplan-Meier curves and time-dependent AUCs; calibration, concordance, decision-curve analysis, and multivariable adjustment against complete clinicopathological covariates were not available. Fourth, the external cohort was modest in size and did not show statistically significant survival separation. Fifth, the fitted gene coefficients and a locked implementation of the score were not available for reproducible deployment. Finally, the computational associations were not tested by targeted molecular experiments, prospective samples, single-cell analysis, or functional perturbation.

Future work should reconstruct the derivation pipeline with explicitly versioned input matrices, probe-to-gene mappings, normalization, batch assessment, and a fully locked coefficient vector. Model evaluation should use nested resampling or a strictly separated feature-selection workflow, include complete clinical covariates, report calibration and concordance with confidence intervals, and test transportability in larger independent cohorts. Experimental studies should prioritize the contrasting *F2RL1*-*TFAP2A* and *FBLN5*-*PLIN5* axes, resolve *HGF* and *FYN* at the protein-activation and cell-compartment levels, and determine whether oxidative-stress perturbation changes the predicted risk-state transcriptional program.

## 5. Conclusions

An integrative analysis of public lung adenocarcinoma transcriptomes identified 44 genes linking tumor-associated differential expression, oxidative-stress annotation, and a redox-associated co-expression module. A seven-gene score based on *FBLN5, HBB, FYN, HGF, TFAP2A, PLIN5*, and *F2RL1* captured survival-associated variation and distinguished proliferative, metabolic, and immune-expression states. The model performed consistently in training and internal validation, while external testing showed moderate discrimination and nonsignificant survival separation. These findings support further study of the signature as a framework for oxidative-stress-associated risk stratification, with prospective validation, complete clinical benchmarking, and experimental confirmation required before translational use.

## Notes

### Competing Interest Statement

The authors have declared no competing interest.

