## supplemental files for "An Oxidative Stress-Associated Seven-Gene Prognostic Signature in Lung Adenocarcinoma: Integrative Transcriptomic Analysis Across Public Cohorts"

1 Division of (Bio) Pharmaceutics, Institute of Zhejiang University-Quzhou, 99 Zheda Road, Quzhou 324000, PR China; (X.Z.); (Y.L.); (X. L.); (Q.X.)
2 College of Chemical and Biological Engineering, Zhejiang University, 866 Yuhangtang Road, Hangzhou 310058, PR China; (X.Z.); (Y.L.); (X. L.)
3 The Quzhou Affiliated Hospital of Wenzhou Medical University, Quzhou People's Hospital, Quzhou 324000, PR China.; (L.Z.)
4 Medical School, Quzhou College of Technology, Quzhou, 324000, PR China; （P.S.）; (Z.L.); (M. C.)
5 College of Jiyang, Zhejiang A & F University, Zhuji, China; (M.C.)
6 College of Pharmacy, Zhejiang Pharmaceutical University, Ningbo, 315100, P.R. China; (S. Z.)

### Supplementary Materials

Supplementary Figures S1-S6 present the pan-cancer expression, univariable overall-survival, and immune-correlation analyses for *F2RL1*, *FBLN5*, *FYN*, *HGF*, *PLIN5*, and *TFAP2A*, respectively. These supplementary analyses are descriptive and emphasize the cancer-type-specific direction of each association.

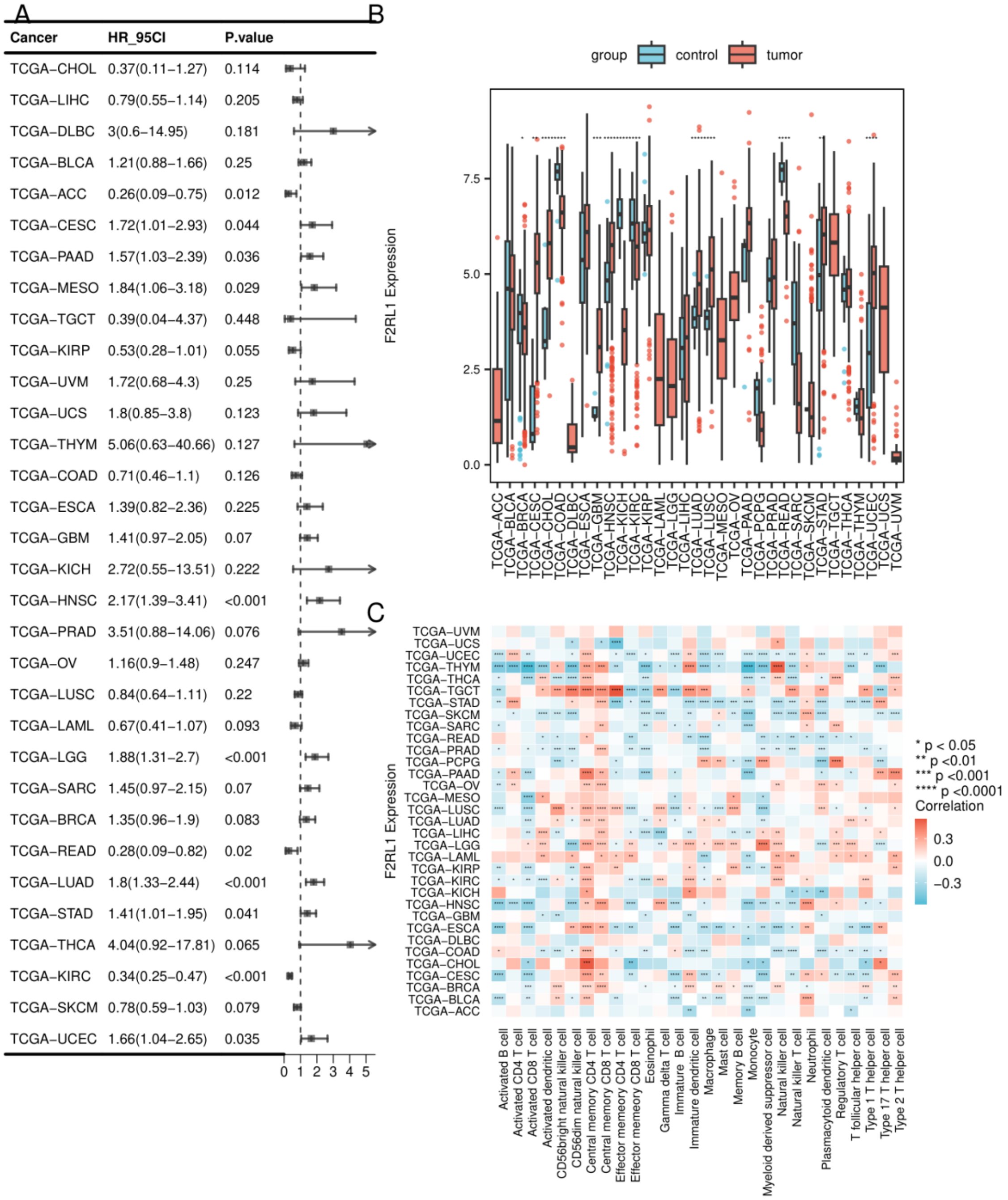


Supplementary Figure S1. Pan-cancer analysis of *F2RL1*. (A) Univariable Cox regression forest plot across TCGA cancer types. (B) Tumor-versus-normal expression distributions across cancers. (C) Correlations between *F2RL1* expression and 28 immune-cell enrichment signatures. Asterisks indicate nominal significance: ****P < 0.0001, ***P < 0.001, **P < 0.01, and *P < 0.05.

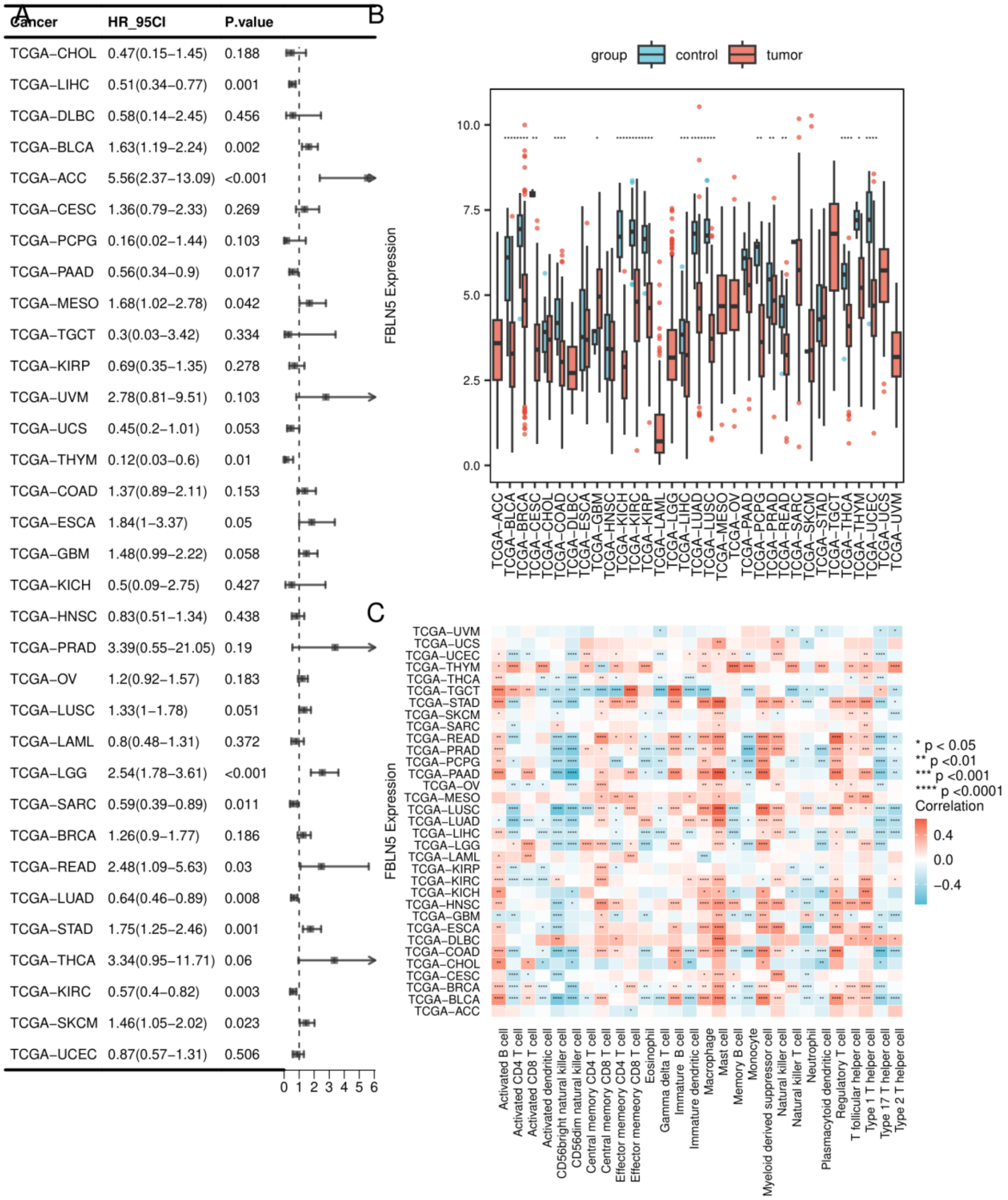


Supplementary Figure S2. Pan-cancer analysis of *FBLN5*. (A) Univariable Cox regression forest plot across TCGA cancer types. (B) Tumor-versus-normal expression distributions across cancers. (C) Correlations between *FBLN5* expression and 28 immune-cell enrichment signatures. Asterisks indicate nominal significance: ****P < 0.0001, ***P < 0.001, **P < 0.01, and *P < 0.05.

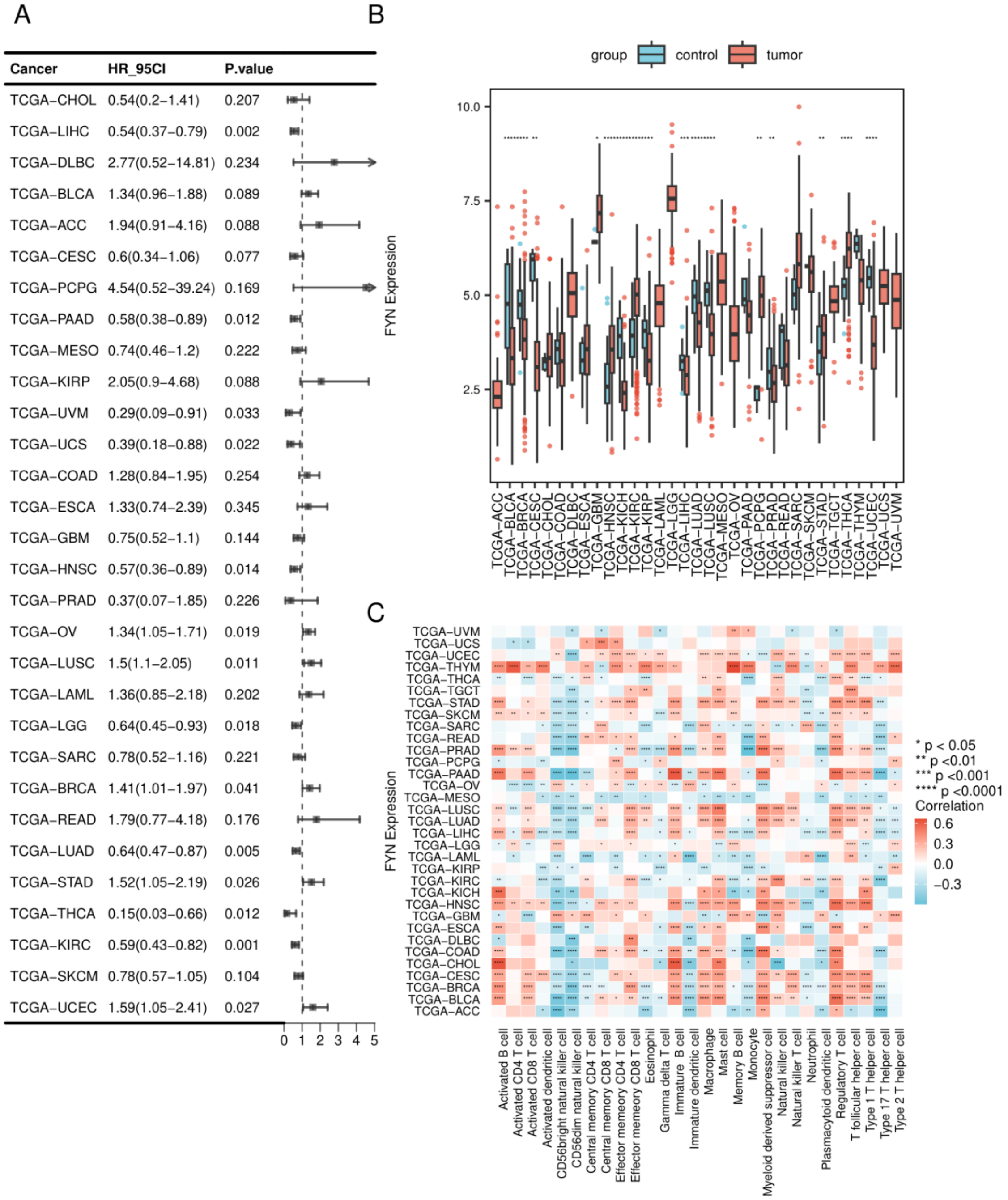


Supplementary Figure S3. Pan-cancer analysis of *FYN*. (A) Univariable Cox regression forest plot across TCGA cancer types. (B) Tumor-versus-normal expression distributions across cancers. (C) Correlations between *FYN* expression and 28 immune-cell enrichment signatures. Asterisks indicate nominal significance: ****P < 0.0001, ***P < 0.001, **P < 0.01, and *P < 0.05.

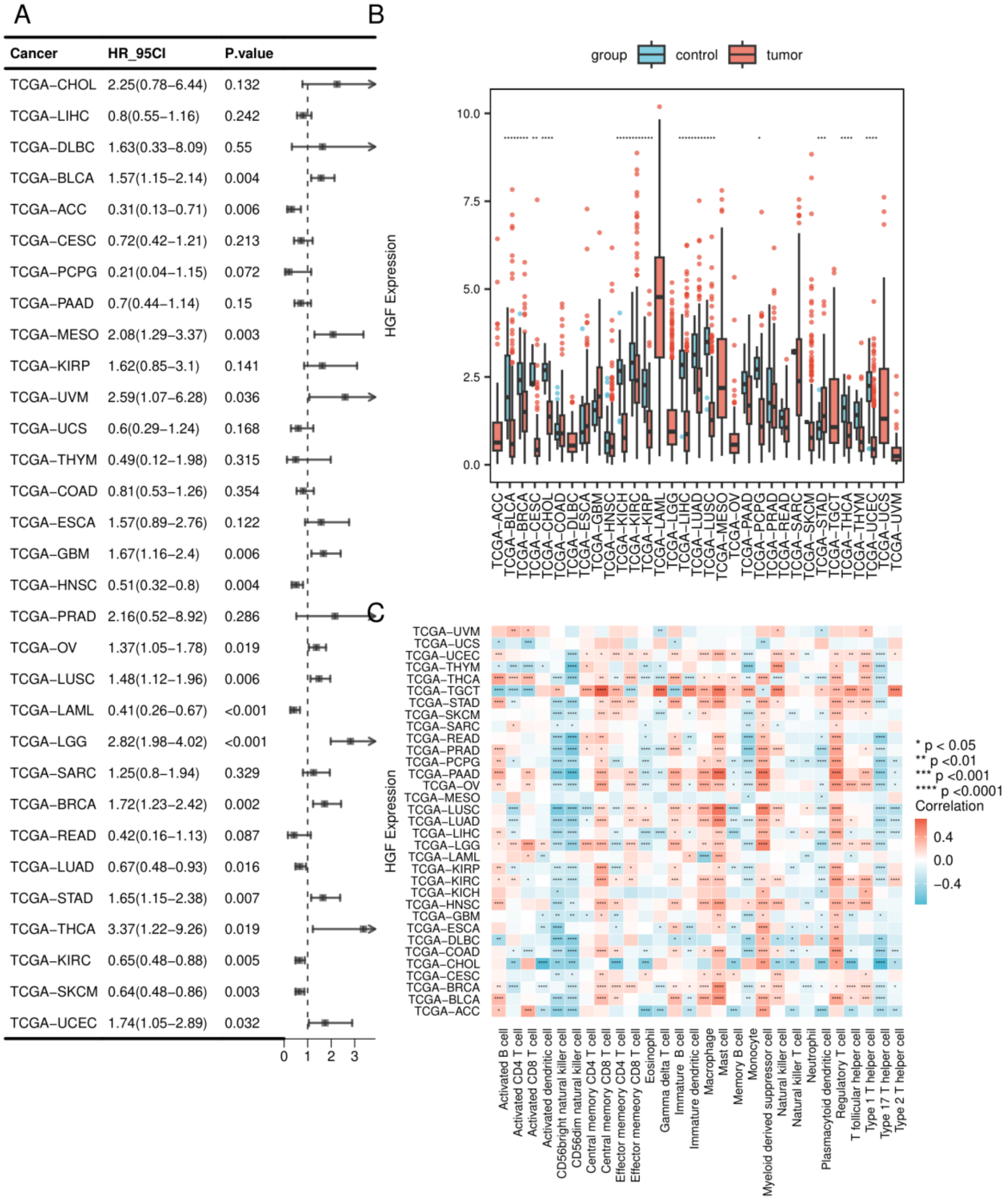


Supplementary Figure S4. Pan-cancer analysis of *HGF*. (A) Univariable Cox regression forest plot across TCGA cancer types. (B) Tumor-versus-normal expression distributions across cancers. (C) Correlations between *HGF* expression and 28 immune-cell enrichment signatures. Asterisks indicate nominal significance: ****P < 0.0001, ***P < 0.001, **P < 0.01, and *P < 0.05.

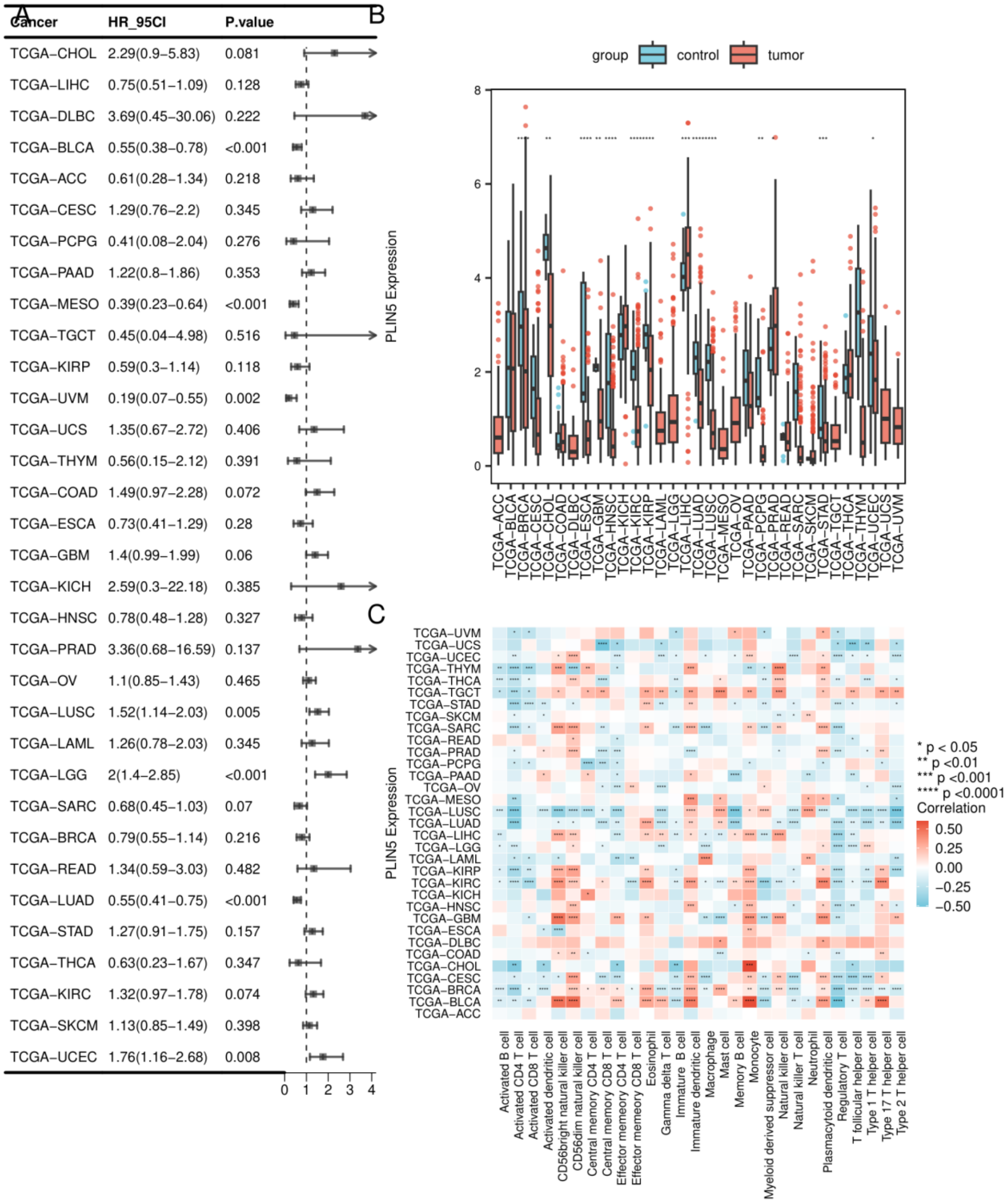


Supplementary Figure S5. Pan-cancer analysis of *PLIN5*. (A) Univariable Cox regression forest plot across TCGA cancer types. (B) Tumor-versus-normal expression distributions across cancers. (C) Correlations between *PLIN5* expression and 28 immune-cell enrichment signatures. Asterisks indicate nominal significance: ****P < 0.0001, ***P < 0.001, **P < 0.01, and *P < 0.05.

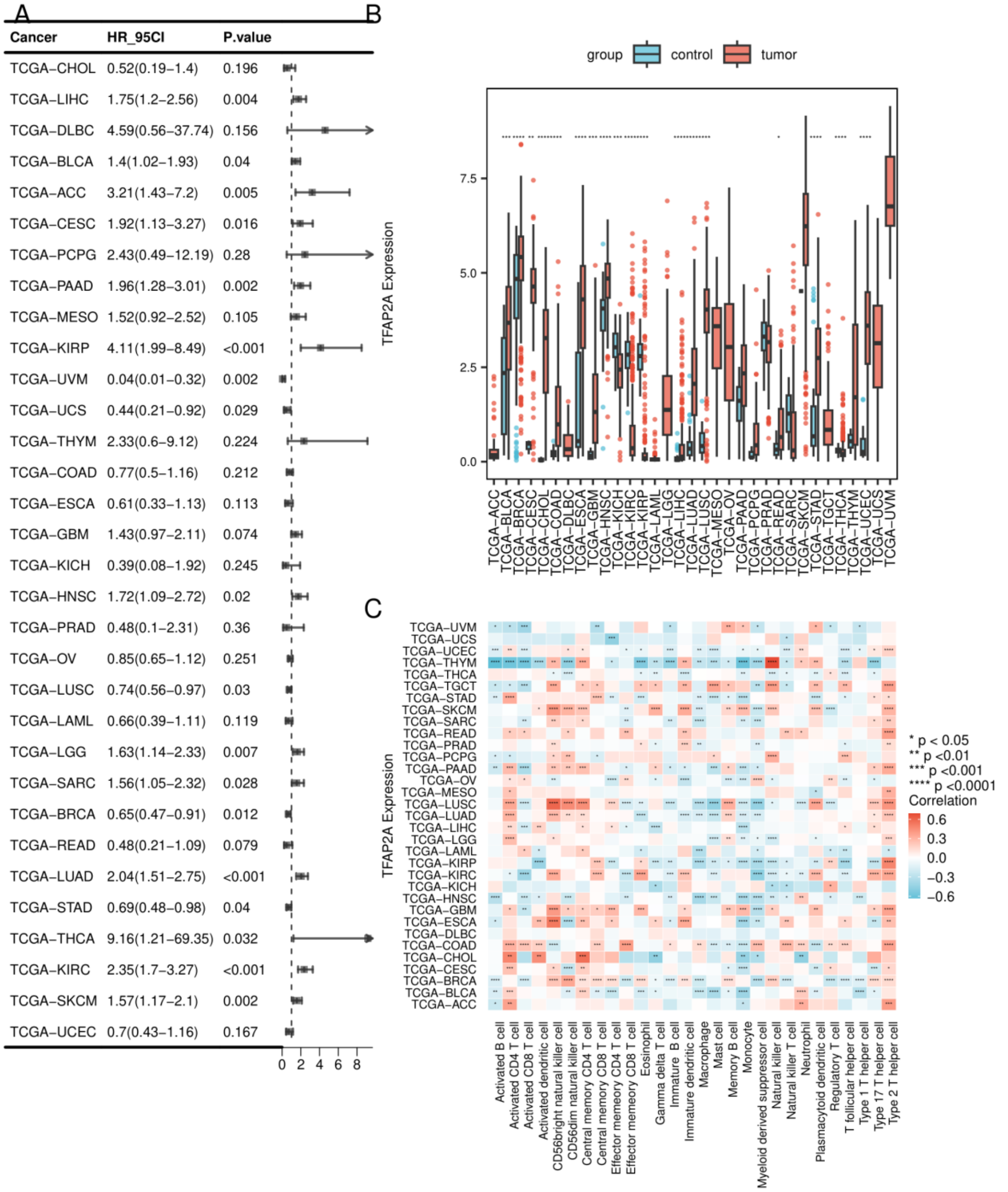


Supplementary Figure S6. Pan-cancer analysis of *TFAP2A*. (A) Univariable Cox regression forest plot across TCGA cancer types. (B) Tumor-versus-normal expression distributions across cancers. (C) Correlations between *TFAP2A* expression and 28 immune-cell enrichment signatures. Asterisks indicate nominal significance: ****P < 0.0001, ***P < 0.001, **P < 0.01, and *P < 0.05.
